# Learning graph networks: sleep targets highly connected global and local nodes for consolidation

**DOI:** 10.1101/2021.08.04.455038

**Authors:** GB Feld, M Bernard, AB Rawson, HJ Spiers

**Author notes:** Gordon Feld, Central Institute for Mental Health, J5, 68159 Mannheim, Germany. These authors contributed equally (GB Feld, M Bernard).

## Abstract

Much of our long-term knowledge is organised in complex networks. Sleep is thought to be critical for abstracting knowledge and enhancing important item memory for long-term retention. Thus, sleep should aid the development of memory for networks and the abstraction of their structure for efficient storage. However, this remains unknown because past sleep studies have focused on discrete items. Here we explored the impact of sleep (night-sleep/day-wake within-subject paradigm) on memory for graph-networks where some items were important due to dense local connections (degree centrality) or, independently, important due to greater global connections (closeness / betweenness centrality). A network of 27 planets (nodes) sparsely interconnected by 36 teleporters (edges) was learned via discrete associations without explicit indication of any network structure. Despite equivalent exposure to all connections in the network, we found that memory for the links between items with high local centrality or high global centrality were better retained after sleep. These results highlight that sleep has the capacity for strengthening both global and local structure from the world and abstracting over multiple experiences to efficiently form internal networks of knowledge.

## Introduction

Sleep has been shown to support the consolidation of declarative memories (Diekelmann & Born, 2010; Feld & Born, 2017; Stickgold, 2005). A night of sleep will tend to enhance memories compared to a similar period of wakefulness during the day (e.g., Abel & Bauml, 2014; Baran, Daniels, & Spencer, 2013; Fenn & Hambrick, 2012). The main mechanism driving this consolidation is thought to rely on the repeated reactivation of recently encoded memories during sleep (Rasch, Buchel, Gais, & Born, 2007; Wilson & McNaughton, 1994). Over time, the reactivation of overlapping information leads to memory abstraction such that some of the detail is lost and the gist of an experience or the centrally important information is retained (Lewis & Durrant, 2011). Sleep is thought to be particularly important for the extraction of such gist and the building of schemas (Lutz, Diekelmann, Hinse-Stern, Born, & Rauss, 2017; Schapiro et al., 2017). Consistent with this, important items encoded before sleep have been shown to be more enhanced by sleep (Feld, Besedovsky, Kaida, Munte, & Born, 2014; Javadi, Tolat, & Spiers, 2015; McNamara, Tejero-Cantero, Trouche, Campo-Urriza, & Dupret, 2014; Wilhelm et al., 2011). In addition, memory strength and item difficulty affect how much memory is boosted by sleep with low strength and high difficulty items profiting the most (Drosopoulos, Schulze, Fischer, & Born, 2007; Kuriyama, Stickgold, & Walker, 2004; Schapiro, McDevitt, Rogers, Mednick, & Norman, 2018).

While most studies of sleep have focused on discrete items such as word lists or item pairs (Diekelmann & Born, 2010; Feld & Born, 2017), most real-world information is interlinked and integrated in networks of knowledge (Patterson, Nestor, & Rogers, 2007). Thus, it remains unclear how sleep impacts the learning of network structures. It has recently been argued that the hippocampus and parahippocampal structures which support spatial memory and navigation may have evolved in humans to support the learning of knowledge networks more broadly (Behrens et al., 2018; Bellmund, Gardenfors, Moser, & Doeller, 2018; Epstein, Patai, Julian, & Spiers, 2017; George et al., 2021; Spiers, 2020; Whittington et al., 2020). This has extended to concepts in reinforcement learning where optimal policies for learning new information need to be developed (Stachenfeld, Botvinick, & Gershman, 2017).

Recordings from individual cells in the hippocampal-parahippocampal network have provided evidence that neurons with specific tunings support aspects of representing a cognitive map of the environment (Epstein et al., 2017; Grieves & Jeffery, 2017; John O’Keefe & Nadel, 1978). Hippocampal place cells in rats show spatially localised patterns of activity during movement through environments with each place cell being active in different specific regions of an environment (O’Keefe & Dostrovsky, 1971). Collectively they provide a unique code for each location encountered in the environment. During periods of sleep and immobility, subpopulations of place cells tend to re-activate, with the order of the cells active ‘replaying’ the sequence of locations visited in an environment previously (Foster, 2017; Ji & Wilson, 2007). Such replay appears to preserve the topological structure of the environment, with sequences of replay along routes in a Y-shaped maze consistent with the physical connections within the environment (Wu & Foster, 2014). This suggests that during sleep, hippocampal networks will replay the various paths experienced during the awake state preserving the structure and may replay intersections or paths that are more frequently encountered if replay is linked to the amount of exposure. For example, passing through a central node many times while exploring a network of paths would lead to reactivations passing through that node many more times than other regions.

Using a film simulation of a complex network of recently learned city streets it has been possible to examine evoked hippocampal responses to street networks when navigating (Javadi et al., 2017). When entering new street junctions, if the new street contained more local streets to choose from (higher degree centrality) then posterior hippocampal activity increased, but if the options decreased (e.g. a dead end) then posterior hippocampal declined. While posterior hippocampal activity responded to local connectivity, the anterior hippocampus responded to changes in the globally connectivity (closeness centrality). Its activity increased when entering a more globally connected street in the network and decreased when entering less globally connected streets. While this suggests the day after learning a street network the hippocampus is able to track the connectivity in a network during navigation, we still know very little about how such networks are learned and consolidated in the gap between learning and navigation.

Recently a number of studies have begun to explore how graph structures may be learned (Lynn & Bassett, 2020; Tomov, Yagati, Kumar, Yang, & Gershman, 2020). However, most of these studies only tracked memory for a short period of time or only investigated learning (Kahn, Karuza, Vettel, & Bassett, 2018; Karuza, Kahn, Thompson-Schill, & Bassett, 2017; Lynn, Kahn, Nyema, & Bassett, 2020; Schapiro, Rogers, Cordova, Turk-Browne, & Botvinick, 2013). In contrast, a recent study asked participants to learn structured information according to a graph and retrieve it 24 hours later in an MRI scanner (Garvert, Dolan, & Behrens, 2017). Neuronal activity measured in the entorhinal cortex tracked the distance between items within the learned graph. Another interesting study (however with a short retention interval ranging minutes) investigated how local connectivity, i.e., community structure, affects statistical learning and could show that participants are sensitive to this type of topology inasmuch as they identified edges connecting local communities as natural breaking points (Schapiro et al., 2013). However, to our knowledge there has been no research on the impact of local and global centrality (i.e., degree centrality and closeness centrality) on information processing and memory acquisition in the long-term. Nor have studies examined how sleep may impact learning networks, where theories emphasize the importance of extracting the gist from experience, which arguably would relate to the centrality of nodes in a network.

Here, we examine how sleep during retention affects associations that were learned using an explicit graph-learning task with a topology that allowed us to disentangle contributions of local and global centrality. We expected 1. that weaker/more difficult associations would be improved more by sleep (as has been demonstrated elsewhere Drosopoulos, Schulze, Fischer, & Born, 2007; Kuriyama, Stickgold, & Walker, 2004; Schapiro, McDevitt, Rogers, Mednick, & Norman, 2018), meaning that greater distance between nodes would predict a greater benefit from sleep, 2. that important information would be improved more by sleep, such that high centrality would predict a greater benefit from sleep. In our design edges connected to high and low centrality nodes did not systematically differ in exposure during learning as we carefully and pseudorandomly chose routes. We did not have strong predictions that sleep more strongly impacts information of high global or local centrality, since relevance has been shown to generally enhance the sleep effect (Wilhelm et al., 2011). In addition, we contrasted centrality derived relevance with more classical reinforcement related relevance, by associating some of the nodes with monetary reward and punishment.

## Methods

### Participants

Twenty-five healthy young men aged between 18 and 30 years (24.20 ± 3.53) took part in the study. Participants were non-smokers, fluent in English, not currently under medication and did not have any physical or mental disorders. They all reported having a regular sleep schedule, going to bed before midnight (11:18 pm ± 47 minutes) and waking up before 8:00 am (7:41 am ± 47 minutes). In addition, participants did not work during night shifts and were not diagnosed with sleep disorders and they did not travel across time zones. Finally, they did not report any stressful events such as exams or deadlines before or during the experiment. Written informed consent was obtained from each participant before starting the experiment. The experiment was approved by the UCL ethics committee (ID number: 8951/002). Participants were compensated financially for their participation.

### Design and procedure

The study was performed in a balanced, within-subject design where participants came for two sessions separated by at least 7 days (8.44 ± 2.98). Each session was composed of a learning and a retrieval phase with a retention interval of 10 hours between the two phases. At the end of the learning phase, participants were asked to avoid learning new information or studying and to not rehearse the information they had learned. Two conditions, sleep and wake, were tested in the experiment for each participant. In the wake condition, participants came at 8 am to complete the learning phase and returned at 8 pm for the retrieval phase. In the sleep condition, participants arrived at 8 pm and returned the following day at 8 am (see Figure 1A). The sequence of conditions was counterbalanced across participants.

**Figure 1.**
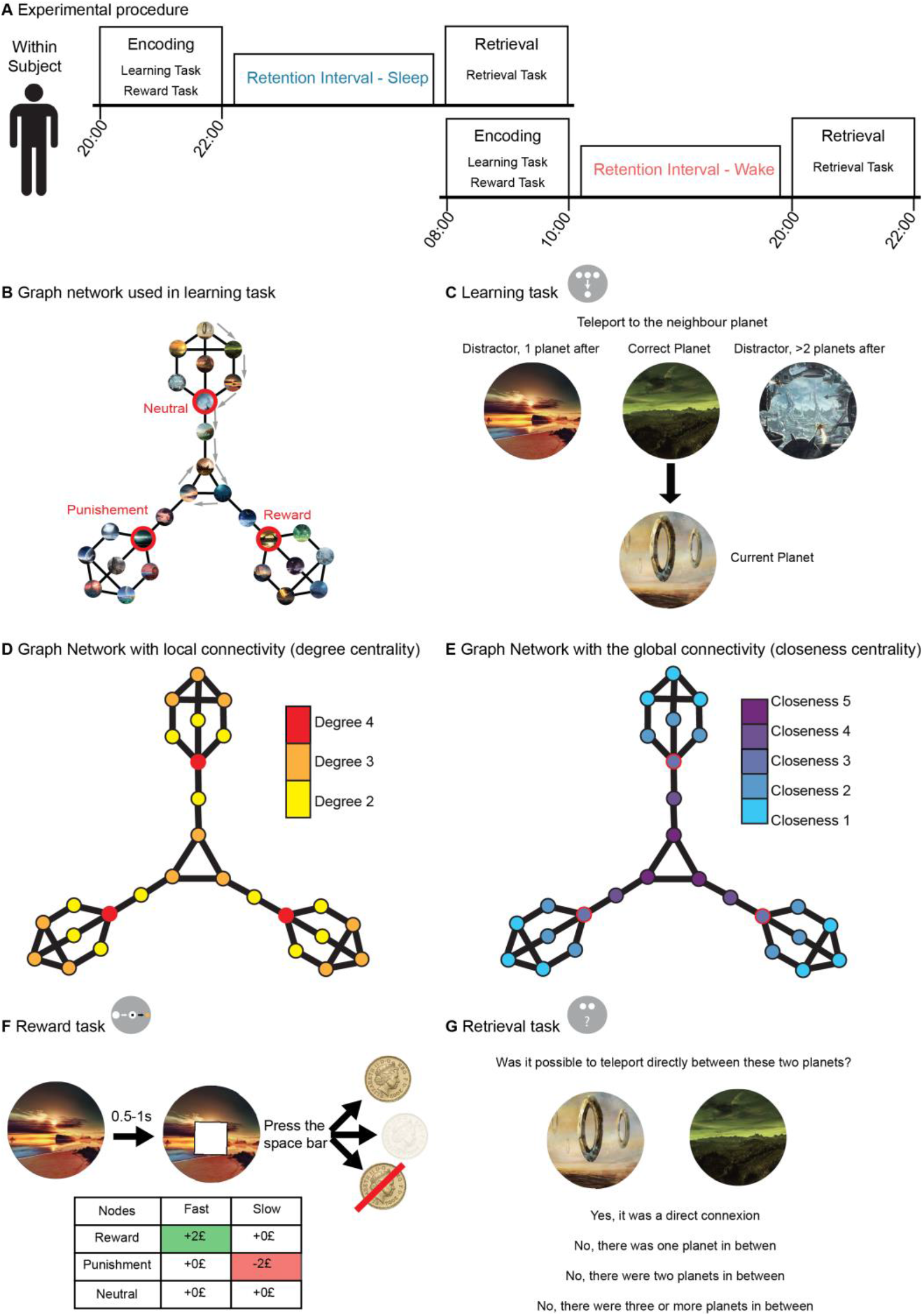
Experimental procedure and task description. A) In our within-subject design participants took part in two identical experimental sessions with parallel versions of the task and retention intervals containing either sleep or wakefulness. The learning phase started at 8:00 am (or pm) and participants performed a learning task (see C) and a reward task (see F). After the 10-hour retention interval, participants came back to the lab to complete the retrieval phase (see G). After one week, participants returned to perform the other experimental session with the remaining retention interval. B) Representation of the undirected graph structure composed of 36 edges (black lines) linking 27 nodes (circles with examples of stimuli presented during the experience). Red nodes represent reinforced nodes, either positive (reward), negative (punishment) or neutral. This image was never shown to the participants, but determined the structure of the learned associations. C) During learning, participants saw one of the three planets displayed at the top. After choosing, the choice was marked but only the correct planet (i.e., the one connected to the bottom planet according to the graph) moved down to replace the bottom planet. Then a new set of three planets appeared at the top prompting a new choice. Participants performed eight such choices (transitions) taking an eight-step route through the graph (an example route is indicated by arrow in B). After each route, they received feedback on their performance and a new route started at a pseudo-random location on the graph. Participants performed 81 routes in total. D) To construct the graph the graph-theoretical parameters of degree centrality and E) closeness centrality were orthogonalized (i.e. allowing to independently asses their effect on retention) and a three-fold symmetry was pursued (to enable equal positions for the reinforced nodes). F) During the reward task, participants were shown a planet representing one of the three reinforced nodes (reward, punishment and neutral). After 0.5-1 seconds a white square appeared on top of the picture. Dependent on the participants pressing the spacebar quickly enough, they were shown the outcomes at the bottom. G) During the retrieval task, participants were shown two planets taken pseudo-randomly from the graph network and had to answer whether they were directly connected or whether one, two or three and more planets were in between.

The experiment was divided into two phases, in the first, the learning phase, participants completed the learning and the reward tasks and in the second, the retrieval phase, they completed the recall task (for details, see task description below). At the end of each phase, control measures were taken. Mood of the participants was assessed by asking them to fill in the Positive And Negative Affective Scale (PANAS, Watson, Clark, & Tellegen, 1988) their subjective sleepiness was measured with the Standford Sleepiness Scale (SSS, Hoddes, Zarcone, Smythe, Phillips, & Dement, 1973) and reaction time (RT) and vigilance were obtained from the 5-minutes version of the Psychomotor Vigilance Task (PVT, Dinges et al., 1997). Additionally, at the end of each retrieval phase, participants performed a word generation task to assess their ability to retrieve highly consolidated information (Aschenbrenner, Tucha, & Lange, 2000).

Finally, at the end of the second retrieval phase, participants filled in the Santa Barbara Sense-Of-Direction scale (SBSOD, Hegarty, Richardson, Montello, Lovelace, & Subbiah, 2002) asking questions about spatial and navigational abilities and completed the Navigational Strategies Questionnaire (NSQ; Brunec et al., 2019) asking questions about their experiences with navigation and their navigation strategy.

### Graph structure

A graph consisting of 27 nodes was constructed (A representation of the graph can be seen in Figure 1B) and pictures were assigned to each node (unique landscapes of extraterrestrial planets). The number of nodes was chosen to enable effective encoding within the 1.5 hours of the learning phase (as determined by pilot participants’ performance). The graph contained 36 edges that connected the nodes. During construction, the graph-theoretical parameters of closeness centrality and degree centrality were orthogonalized and a three-fold symmetry was pursued (Figure 1D and 1E). In addition, three nodes corresponding to the red nodes in Figure 1B were selected to be reinforced. The nodes were either positively reinforced (reward node), negatively reinforced (punishment node) or not reinforced (neutral node). The reinforcements were associated during the reward task (for details, see task description below). This graph was never shown to the participants during the experiment and participants were not explicitly told about an underlying structure of the learning task.

### Learning task

The learning task was gamified to optimise participants’ motivation during learning and retrieval (see Supplementary Methods for details). Briefly, the task was embedded in a storyline of humankind on the brink of extinction on earth and participants explored planets to find a new home for humans to live. Our piloting demonstrated that participants’ motivation, especially during the 1.5 hours of the learning session, greatly benefitted from this approach, which enabled us to use a somewhat larger graph. To familiarise the participant with the stimuli, each planet was shown in the middle of the screen with its name under it for 2 seconds with an inter-stimulus-interval of 0.5 seconds. Next, the participants learnt the graph structure by performing 81 routes of 8 transitions length between the planets of the graph. An overview of the task can be seen in Figure 1. For each transition, the participants were asked to identify the neighbour of the current planet that was presented at the bottom of the screen (i.e., the planet connected by a single edge) while being shown three potential planets at the top of the screen. One option was the correct planet (one of the 2-4 connected planets) and the two other options were incorrect (i.e., not directly connected). Of the two wrong choices, one of the planets had a distance of two edges, i.e. there was one planet between the current planet and the incorrect choice, and the other had a distance of three or more edges, i.e., there were at least two planets in between. The wrong choices were chosen randomly from all planets qualifying the distance argument and only the shortest distance was considered relevant for this choice. During each transition, participants had a maximum of 10 seconds to choose the correct transition. If they did not answer within the time limit, the choice was considered incorrect and the next trial was presented.

### Reward task

The reward task was constructed to be an adaptation of the monetary incentive delay task that robustly activates reward areas (Knutson, Westdorp, Kaiser, & Hommer, 2000). During the task, participants saw the three reinforced nodes mentioned above. During 180 trials, a fixation cross appeared in the middle of the screen for 250 milliseconds followed by one of the three planets representing the nodes for 2 seconds. After 0.5 to 1 seconds, a white square appeared and the participants needed to press the spacebar as fast as possible (pressing before the square was shown was considered a miss). Depending on their RT and the planet presented, the participants obtained a different monetary outcome (Figure 1F). For the reward planet, if the participants were fast enough, they got +2£, but if they were not fast enough, they got 0£. For the punishment planet, being fast enough let them earn 0£, but being too slow made them lose −2£. Finally, for the neutral planet participants got 0£ whether they were fast or not. Their wins and losses accumulated resembling the amount they would receive for performing the reward task.

### Retrieval task

The retrieval task was divided into three parts. In the first part, we presented two planets (without their names) next to each other to the participants that were chosen pseudo-randomly and asked if they were directly connected during learning or if there were one, two or three and more planets in between (Figure 1G). This was done for all the possible combinations of the 27 planets, therefore the participants were presented with 351 trials. In the second part, we asked the participants about the names of the planets. Participants performed 27 trials, one for each planet, and were asked which of four possible names was correct. The three incorrect names were chosen to be from planets that were one, two or more edges away, respectively. In the last part, the participant identified the contingencies learned in the reward task again.

### Data reduction and statistical analysis

The data of 6 participants were excluded from the analysis. Four participants had a learning performance with an accuracy lower than 0.5 and two participants had a high learning performance, accuracy above 0.8, but a low retrieval performance, less than 0.4. In addition, only node pairs with distance 1 to 4 were analysed since the retrieval tested participant’s knowledge of the graph structure only to distance 4 at maximum. Data reduction was performed in Matlab 2018a and the statistical analysis depended on R studio (Version 1.0.143). The analysis relied mainly on repeated measures ANOVA, paired t-tests, pearson correlations and regression with linear modelling. Details regarding the data reduction and statistical analysis can be found in the supplementary methods.

## Results

### Retention performance

A retention measure was created by subtracting the learning performance from the retrieval results, i.e., how much information the participant retained during the retention period (Figure 2A; for more details see the Supplementary Methods section and Supplementary Figure 4). When collapsing all distance information, we found that participants retained more information from learning to retrieval in the sleep condition than in the wake condition (t(18)=2.13; p<0.05). An ANOVA across distances, confirmed this main effect of interval (F(1,18)=4.52; p<0.05). When analysing the distances individually we found that for distance 4 the sleep condition also showed better retention (t(18)=2.29; p<0.05) (Figure 2D). However, no such difference was found for distances 1, 2 and 3 (p>0.19). Using linear regression lines modelling the distances per interval revealed a difference of intercept between sleep and wake conditions (F(1,18)=-3.99; p=0.02). However, this analysis did not reveal a difference in the slopes, i.e., no distance by interval interaction (F(1,18)=-2.22; p=0.09, Figure 2G).

**Figure 2.**
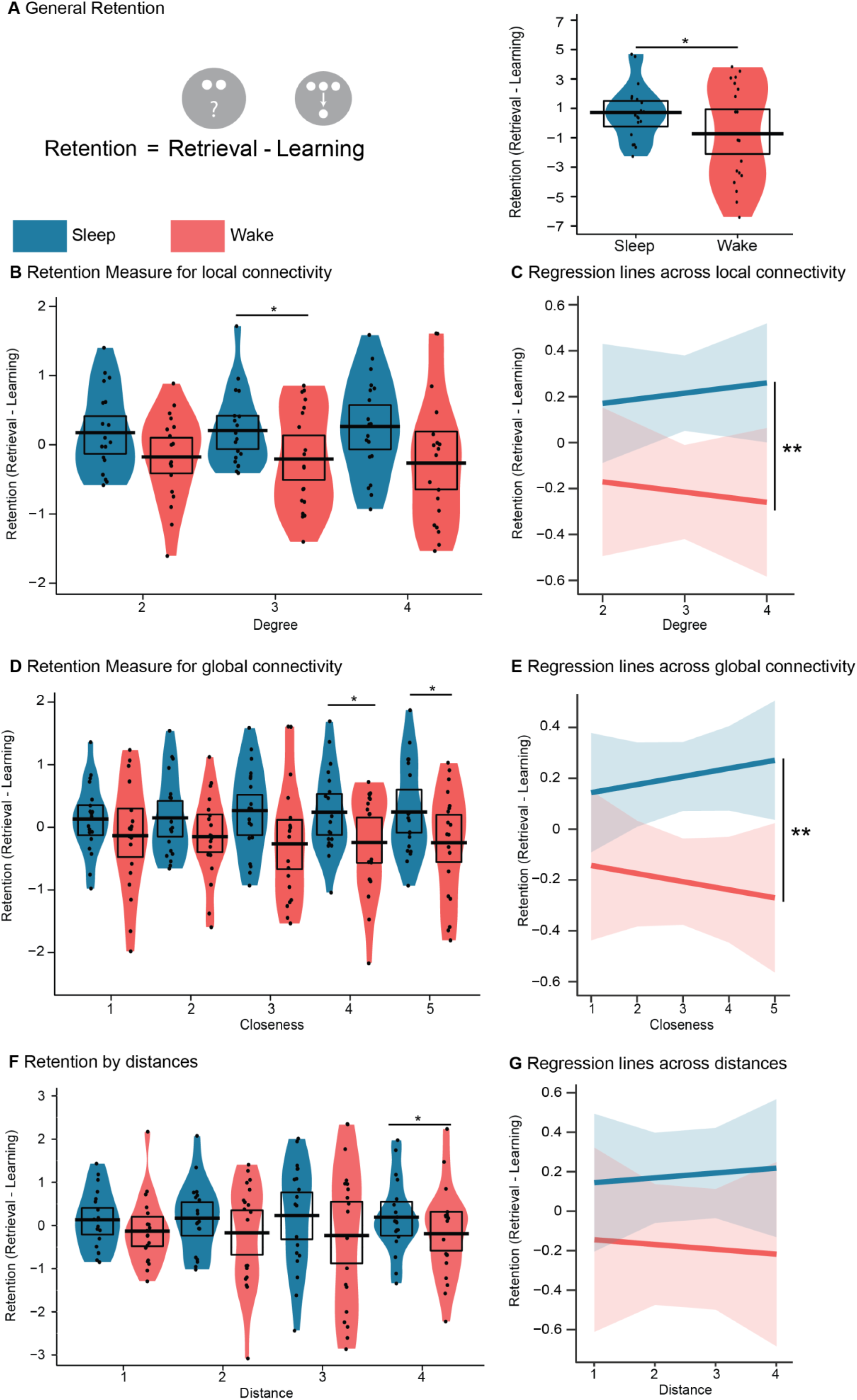
Retention performance. A) Left, the retention measure was calculated by subtracting the learning performance from retrieval performance. Details can be found in Supplementary Figure 5. Right, mean overall retention performance for the sleep (blue) and the wake (red) condition. B) Mean retention performance and C) regression model for the different distances within the graph, D) & E) for the different levels of degree centrality of the nodes and F) & G) for the different levels of closeness centrality of the nodes. For the violin plots, the black dots represent the individual performance, the black bar represents the mean across participants, the black rectangle shows the 95% of a Bayesian highest density interval and the coloured shape displays the smoothed density. *p<0.05, **p<0.01

Regarding the local connectivity (degree centrality), participants retained more information during the sleep interval (F(1,18)=4.71; p=0.04). This effect was stronger for nodes of degree centrality 3 (t(18)=2.15; p<0.05) compared to degree 2 (p=0.06) or 4 (p=0.07, Figure 2B). The regression analysis analysis confirmed the main effect of sleep (difference in intercept: F(1,18)=-4.86; p=0.04) and indicated that higher degree centrality was associated with an increased benefit from sleeping during retention (difference in slope: F(1,18)=-8.26; p=0.01, Figure 2C).

Similar results were found for global connectivity (closeness centrality) since participants again performed better across the sleep retention interval (F(1,18)=5.26; p=0.03), which was mirrored by a sleep benefit for closeness centrality of 4 (t(18)=2.27; p=0.04) and 5 (t(18)=2.90; p=0.001) but not for closeness 1 (p=0.28), 2 (p=0.17) or 3 (p=0.07) (Figure 2D). Again the regression analysis showed that participants performed better across sleep (difference in intercepts: F(1, 18)=-3.79; p<0.01) and that a higher degree centrality increased the effect of sleep (difference in slopes: F(1, 18)=-3.59; p=0.01, Figure 2E).

### Learning and retrieval performance

For the learning results, there was a trend towards overall learning performance (calculated by averaging the encoding results over the four distances, see supplementary methods) being greater for the wake than for the sleep condition (t(18)=-2.09; p=0.06) (Supplementary Figure 1B). However, when viewing only first order connections (as learned during the task) and dividing the learning task into thirds, no main effect of sleep or wake was evident when analysing the three thirds in an ANOVA (F(1,18)=2.66; p=0.12) or for the last third in an individual t-test (t(18)=-1.09; p=0.29, Supplementary Figure 1A). Although the participants increased their learning performance across the thirds (F(1, 18)=124.74; p<0.001). The individual learning curves of participants are shown in Figure 3 A and B for both conditions.

**Figure 3.**
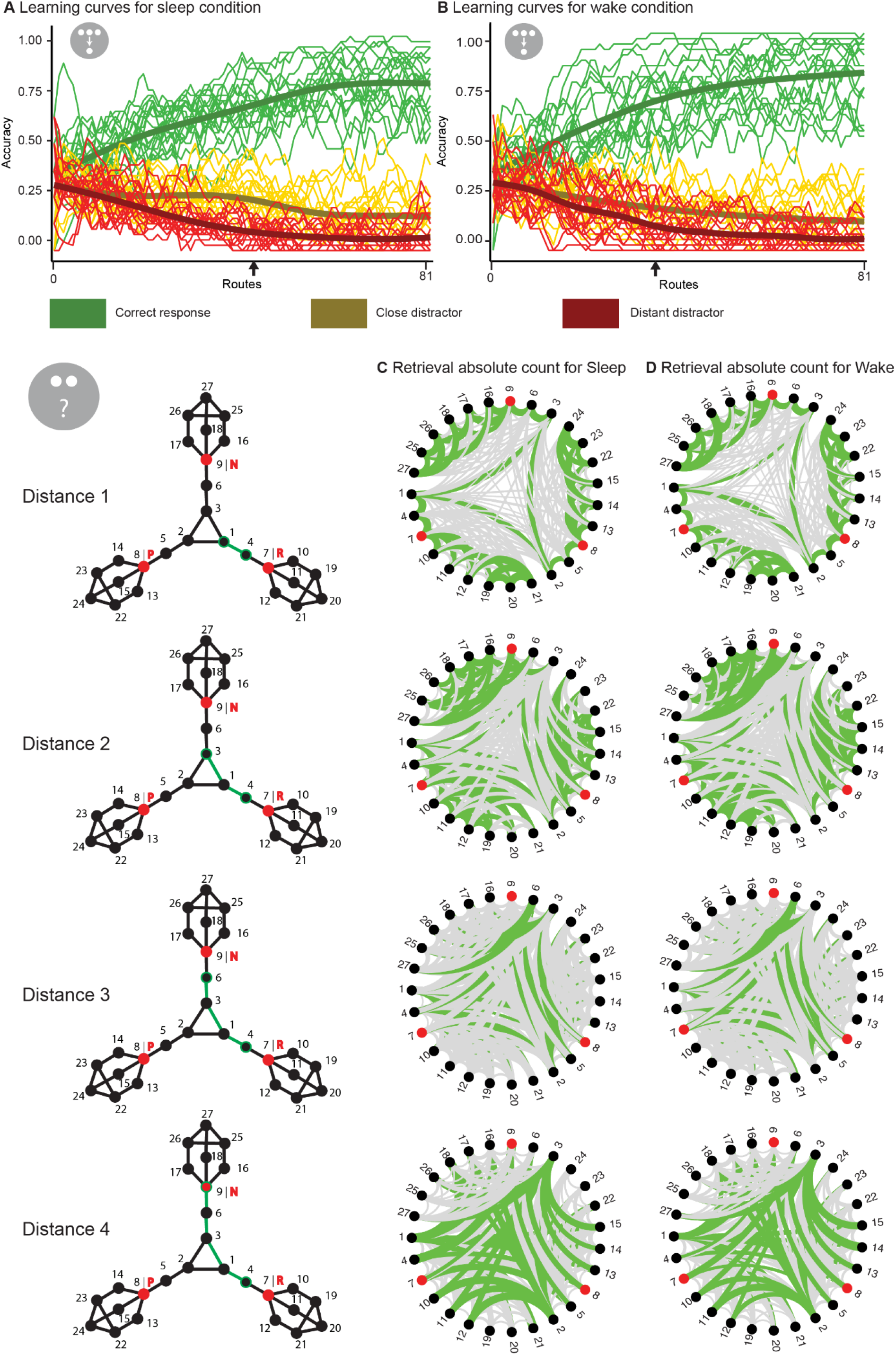
Participants’ raw learning and retrieval data. A) Learning curve across the 81 routes for the sleep and B) the wake condition. Mean (thick line) and individual responses (thin lines) for correct response (green), the close distractor (yellow) and the distant distractor (red) smoothed by a five point moving average. The black arrow indicates when participants performance was significantly biased by the graph structure (i.e. accuracy of the close and distant distractor started to differ).C) Retrieval task data for each distance (rows) in the sleep and D) the wake condition. On the left the graph structure and an example (in green) is shown for each distance, which is defined by the number of edges between the pair of nodes tested. Within the circles, green lines represent correct connections and grey lines correspond to incorrect connections at that distance, whereas line thickness depicts how many participants gave the respective answer.

Regarding local connectivity, an ANOVA showed that higher degree centrality was associated with better learning (F(1, 18)=44.55; p<0.001) but no effect of intervals nor an interaction was found (p>0.5) (Supplementary Figure 2A). For global connectivity, similar results were found, i.e., participants learned nodes with higher closeness centrality better (F(1, 18)=33.01; p<0.001) (Supplementary Figure 2C) but there was no effect for the retention interval and no interaction (p>0.5). Similarly, no difference in intercept and slopes were found for the linear model analysis.

Regarding the general retrieval performance (calculated by the hit rates for each distance, see supplementary methods), participants were better at correctly identifying closer pairs than more distant ones (F(1,18)=7.46; p<0.001) but there was no main effect of the interval nor an interaction effect (p>0.5) (Supplementary Figure 1C). A visualisation of the raw retrieval data can also be found in Figure 3 C and D. No statistical difference was found for local connectivity (Supplementary Figure 2B) but global connectivity showed a main effect of the centrality (F(1, 18)=4.34; p<0.01) (Supplementary Figure 2D). Similar to learning, no statistical differences were found for the linear model analysis. Finally, participants were not able to remember planet name associations better across the sleep than the wake condition (t(18)=0.89; p=0.38)

### Reinforcement

Individual balance curves (the amount of money given to participants, see Methods section), for the two conditions can be found in Figure 4 A/B. Regarding the retention measure, an ANOVA found no influence of sleep or wake (F(1, 18)=2.33; p<0.14) nor was there an effect of reinforcement or an interaction of the two (F(1,18)=1.12; p=0.34) (Figure 4C). However, exploratory paired t-tests found that for the punishment node participants performed better in the sleep condition (t(18)= 2.10; p<0.05) but not for the reward and neutral nodes (p>0.30).

**Figure 4.**
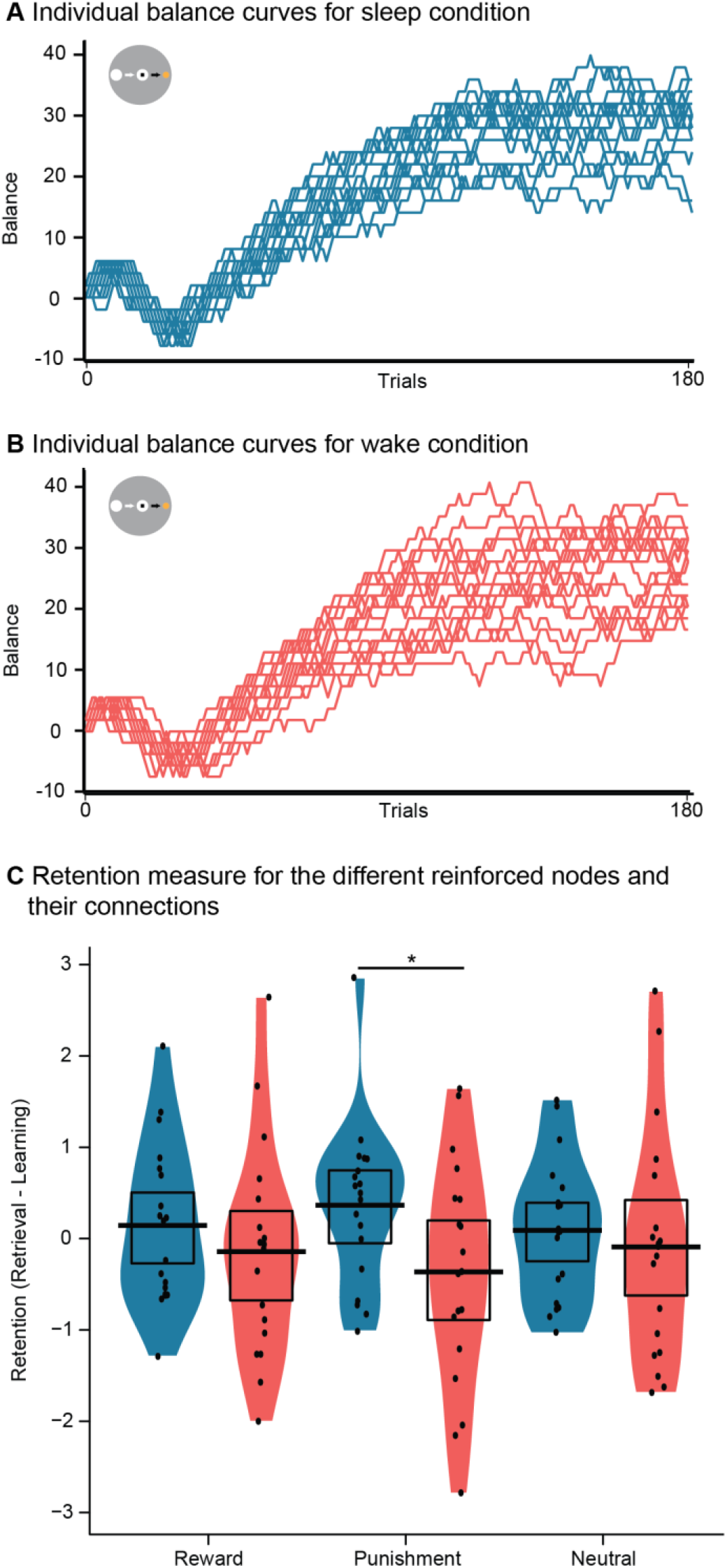
Reinforced nodes results. A) Individual monetary balance curves during the reward task in the sleep and B) the wake condition. Participants earned the amount reached at the end of the task. C) Retention performance for the reinforced nodes. The black dots, bar and rectangle represent the individual performances, the mean and the 95% of a Bayesian highest density interval, respectively. The coloured shape around shows the smoothed density. *p<0.05

### Navigation tests

A significant correlation was found between the two navigation questionaires (r = 0.55, p<0.02). Since the variance of the NSQ was larger further correlations used this navigation test (Figure 5). Participants with a higher score in the NSQ have a higher mapping strategy. We found a significant negative relationship between the NSQ and the retention measure for the sleep (r=-0.58, p<0.01) but not for wake condition (r=-0.40, p>0.09). A positive relationship between the NSQ and learning performance in the sleep condition (R=0.46, p<0.05) and between the NSQ and retrieval performance in the wake condition (R=0.48, p<0.05). Correlations for learning performance in the wake condition (R=0.43, p=0.07) and retrieval performance in the sleep condition (R=0.37, p=0.12) did not reach significance. In general, although some relationships did not reach significance in the sleep or the wake conditions the overall pattern of effects was similar.

**Figure 5.**
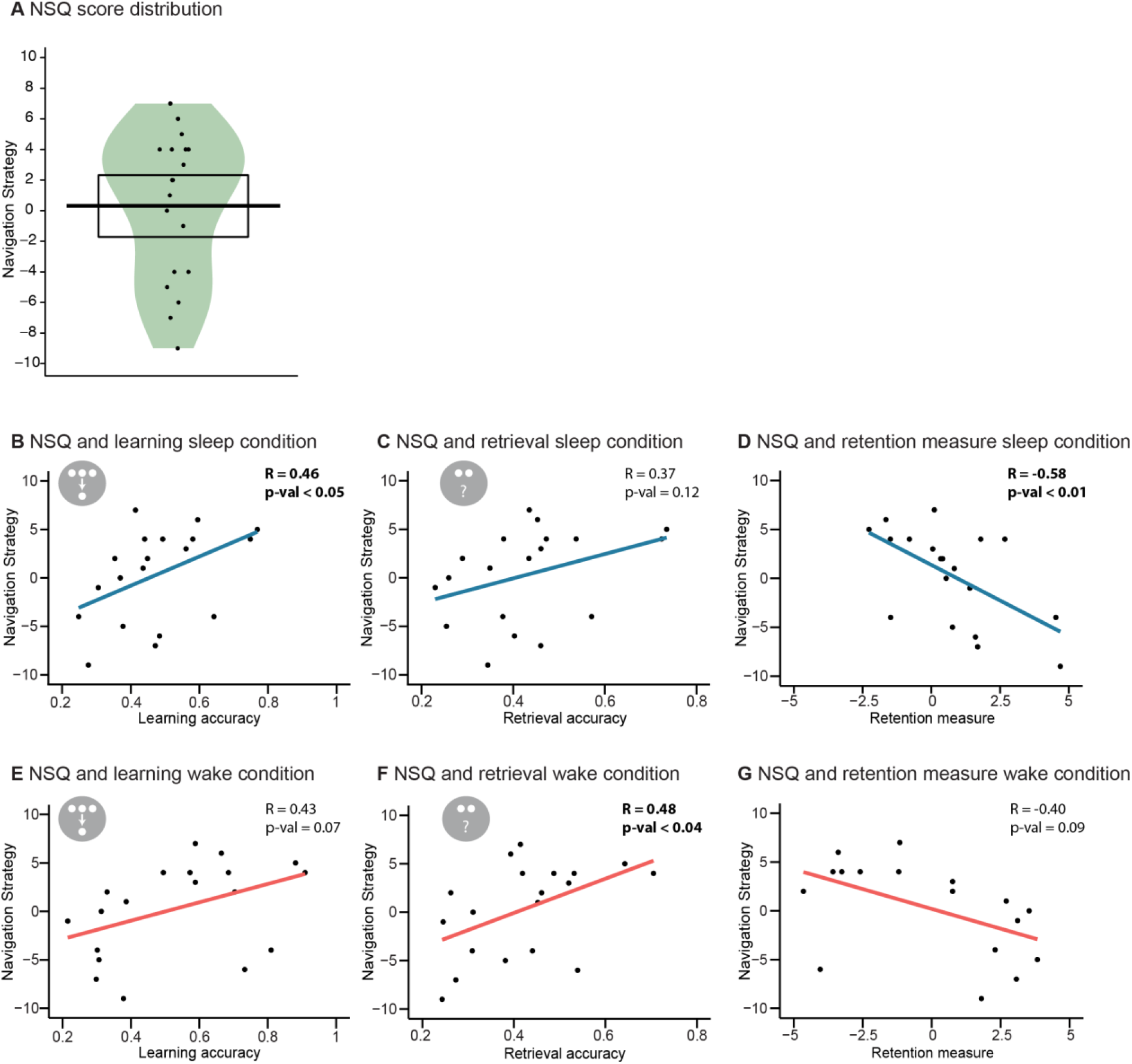
Correlation of the results and the navigation score. A) The Navigation Strategies Questionnaire (NSQ) score, the black dots, bar and rectangle represent the individual performances, the mean and the 95% of a Bayesian highest density interval, respectively. The coloured shape around shows the smoothed density. B) Relationship of the Navigation Strategies Questionnaire (NSQ) with learning performance, C) retrieval performance and D) retention performance for the sleep condition and with E) learning performance, F) retrieval performance and G) retention performance for the wake condition. Regression lines (blue – sleep, red – wake) and black dots for the individual data points are shown.

### Control tasks

There was no difference in the long-term retrieval performance (measured with the word generation task) between the sleep and the wake conditions (t(18)=-0.82; p=0.42). Also, no statistical difference was found for objective vigilance (reaction speed of the PVT), subjective sleepiness (measured by the SSS) or positive or negative affect (measured by PANAS) between the sleep and the wake conditions during learning (PVT: t(18)=1.02, p= 0.32; SSS: t(18)=-0.57, p=0.57; PANAS-positive: t(18)=1.37; p= 0.19; PANAS-negative: t(18)=2.08; p>0.05) or retrieval (PVT: t(18)=0.66, p=0.52; SSS: t(18)=-0.86; p=0.40; PANAS-positive: t(18)=0.67; p= 0.51; PANAS-negative: t(18)=-0.33; p=0.74). Descriptive statistics can be found in Table 1.

**Table 1.** Some data.

|  | **Sleep** |  | **Wake** |  |
| --- | --- | --- | --- | --- |
|  | Mean | SD | Mean | SD |
| *Word Generation* | 41.16 | 2.73 | 42.79 | 2.95 |
| *PANAS - Positive* |  |  |  |  |
| Learning Phase | 30.16 | 1.52 | 28.36 | 2.21 |
| Retrieval Phase | 28.17 | 1.86 | 26.89 | 1.52 |
| *PANAS - Negative* |  |  |  |  |
| Learning Phase | 14.10 | 0.77 | 12.58 | 0.58 |
| Retrieval Phase | 11.84 | 0.59 | 12.00 | 0.55 |
| *PVT* |  |  |  |  |
| Learning Phase | 3.13 | 0.08 | 3.08 | 0.07 |
| Retrieval Phase | 3.17 | 0.08 | 3.14 | 0.06 |
| *SSS* |  |  |  |  |
| Learning Phase | 2.63 | 0.23 | 2.79 | 0.29 |
| Retrieval Phase | 2.42 | 0.18 | 2.69 | 0.22 |

## Discussion

Here, we investigated the impact of sleep on consolidating learned topological networks, which varied across them in global and local connectivity of nodes. We found both connections to globally and locally highly relevant nodes were preferentially enhanced by sleep. This was despite equal exposure to all the connections in the network during learning. By contrast, sleep had no impact on the enhancement of nodes made salient by monetary reinforcement. We discuss how these results help advance our understanding of how representations of learned graph networks are affected by offline processing, models of sleep and consolidation and implications for understanding offline replay in hippocampal networks.

The current study presents a novel associative learning task, where information was learned according to a graph network. We found that during learning, the graph influenced behaviour beyond first level associations inasmuch as close distractors (distractors that were only one edge away from being a direct connection) were more frequently chosen than distant distractors (distractors that were at least two edges away from being a direct connection) when participants made errors. Although sleep enhanced memory retention per se, its effect was not enhanced by increased distance between the nodes, as might have been expected due to sleep preferentially enhancing items with low association and high difficulty (Drosopoulos et al., 2007; Kuriyama et al., 2004; Schapiro et al., 2018). Sleep did however specifically enhance items that were more relevant for navigating the network as items of high local and of high global connectivity showed a stronger sleep effect. This graded effect of topological relevance may be related to findings of graded reward effects on memory within a maze (Braun, Wimmer, & Shohamy, 2018). In this study, participants explored a maze by uncovering cards laid out in a 2-d grid and received a high or a low reward after a certain amount of cards. The reward effect was higher the closer a card was to the final card that was uncovered. Although we found a similar graded effect of topological relevance, we found no effect of monetary reinforcement applied to a subset of nodes. This may likewise be explained by a spread of reward across the network, if one assumes that our network was too small. We chose the size of our network after extensive piloting so that it could be learned to about 80% correct within 1.5 hours. Using a larger network or maybe even two networks with different reinforcement procedures may prove more fruitful.

Sleep has been suggested to enhance the abstraction of gist from episodes (Lewis & Durrant, 2011). Such gist abstraction may become more important, when larger networks are learned, as participants will struggle to keep all connections in memory. Representing the network at different scales would then allow for more efficient memory processing. Integrating very large networks may occur over several nights of sleep, as has been shown for other gist abstraction processes (Lutz et al., 2017). Once a large network has been built, it may prove that new items can be added to this schema much faster and with less reliance on the hippocampus (Tse et al., 2007; van Kesteren, Fernandez, Norris, & Hermans, 2010). In fact, sleep may no longer be required to consolidate such memories since encoding may circumvent the hippocampus directly integrating into the neocortical knowledge network (Himmer, Muller, Gais, & Schonauer, 2017). Based on past work showing this rapid learning of new nodes to networks (Tse et al., 2007; van Kesteren et al., 2010), we would predict that after adding new nodes to our network encoding would not require sleep consolidation to stabilize traces as long as the global topological is minimally affected. This could be studied in our paradigm by adding nodes that do or do not strongly influence topology. For example, providing a single shortcut between very distant parts of the network could radically change the closeness centrality of nodes and potentially drive more extended consolidation during sleep.

One prediction from theories highlighting the importance of global gist extraction and schema development (Gilboa & Marlatte, 2017; Lewis & Durrant, 2011) is that predominantly the globally important information would be prioritised over the local important information. We did not find this was the case, since high degree centrality also impacted consolidation during sleep. In future research, it would be interesting to explore memory after several days to observe whether global and local information is lost or retained at the same rate. Testing other network structures would also help explore whether the local and global effects are additive, in that are nodes with both high local and global centrality doubly enhanced by sleep.

According to one perspective, hippocampal replay during sleep has been considered to be closely aligned with prior experience, so that sequences of place cells evident during learning would emerge again during subsequent sleep (Ji & Wilson, 2007). Reward has been shown to influence replay of place cells and enhancing reward increases the frequency of replay events in brief rest intervals during learning (Ambrose, Pfeiffer, & Foster, 2016). Another study showed that enhancing dopaminergic modulation within the hippocampus enhances replay frequency during post encoding rest (McNamara et al., 2014). An alternative perspective has recently suggested that in the absence of reward replay events may represent random samples from available trajectories through space (Stella, Baracskay, O’Neill, & Csicsvari, 2019). The enhancement of globally relevant nodes in our network could be explained by either account. Either global relevance was inferred already during wake encoding and enhanced replay of those nodes during sleep by a synaptic tagging mechanism (Redondo & Morris, 2011) or the structure of the network may have biased replay that occurs in form of random walks on the graph to emphasize nodes with high betweenness centrality. As betweenness and closeness centrality were highly correlated in our graph, we cannot at present distinguish which the two metrics might impact consolidation during sleep. With a larger network it would be possible to dissociate closeness centrality and betweenness centrality (see Fig 1 of Javadi et al., 2017). If replay takes a random walk through the network it might specifically enhance regions of high betweenness centrality but not closeness centrality. However, it may be that replay prioritizes important structures to be learned (Mattar & Daw, 2018), which recent evidence supports (Liu, Mattar, Behrens, Daw, & Dolan, 2021).

In conclusion, we find that local and global aspects of connections between individual items of a declarative associative memory task determine access to sleep dependent memory consolidation. This approach has the potential to explore in more detail how replay influences the knowledge structure of declarative memory. Many psychiatric disorders come with impaired memory as well as sleep disruption, better understanding how complex memories are formed during sleep may increase our understanding of these disorders.

## Acknowledgements

This work was supported by grants from the German Research Foundation to GBF (DFG; FE 1617/1-1 and FE 1617/2-1) and by a James McDonnell Scholar Award grant to HJS.

## Box 1. Graphs.

We refer to graphs in their mathematical form to describe connections (edges) between instances (nodes) and not, as more commonly used, as a form of data presentation. Graphs are part of our everyday life and are most easily visualized as networks of connections. For example, the connections of the London Underground can be described using a graph. Here, the different stations are the nodes and the connections are the edges, i.e., the node Russel Square is connected to the node Euston via the node King’s Cross and two edges. The degree centrality and closeness centrality of a given node signify its relevance in the network. For example Oxford Circus has a high degree centrality as it is directly connected to 6 other stations, but it also has a high closeness centrality as you can travel to any other node relatively quickly (i.e., using few edges).

## Supplementary Methods

### General procedures

All the learning tasks were programmed and executed on Matlab2016a with the package Psychtoolbox-3. Matlab was running on Dell OPTIPLEX 990 with Windows 7 Enterprise with a Dell P2011H monitor (resolution of 1600×900). Up to four participants were run at one time in individual cubicles.

Two storylines were created and were told using texts during the instructions, as well as, videos before and after the tasks, to keep the participants interested and focused. The first storyline, corresponding to the first session, emphasized that Earth will be destroyed and participants are sent into a new galaxy to explore it. The second storyline, related to the second session, told that there are strange sounds and radio wave frequencies that are being emitted from a neighbouring galaxy. This time, participants must explore this galaxy to lead to further understanding of what these sounds may mean. Videos that helped build this storyline can be found in the supplementary methods. The stimuli were presented on a screen that contained elements of a videogame to further gamify the experience (see below for details).

Both sessions used the same protocol and participants learnt the same graph structure but with a different set of stimuli (i.e. pictures) for each session. In addition, each participant received a uniquely randomised version of the task with different stimuli, planets, names and routes. For each memory task, instructions and examples were presented at the beginning and participants had the opportunity to ask any questions after the instructions.

### Learning task

After each route that consisted of 8 transitions, the participant received feedback of their performance, i.e., they were shown up to three stars with a written sentiment (0 stars – Try harder, 1 star – Well done, 2 stars – Superb, 3 stars – Amazing). Furthermore, participants could keep track of their progress by consulting a bar at the right side of the screen that was updated after each transition. Moreover, a rank system was created and participants were promoted after completing every 27 routes (‘Novice’; ‘Lieutenant Jr.’; ‘Lieutenant’; ‘Captain’; ‘Major’; ‘Colonel’; ‘Commander’). An “energy level monitor” at the bottom was filled up to 1500 at the beginning of the experiment. For additional motivation, the participants lost 2 units each time they got an incorrect answer whereas only 1 unit was lost for a correct choice. This bar was related to the storyline as depleting too much energy would produce negative consequences for humankind.

### Reward task

To learn these associations between planets and gain/loss, the criterion, i.e., the maximum time for the reaction that participants needed to match, changed throughout the task. At the beginning, the criterion was slow so participants would accumulate 6£, even if they were reacting rather slowly, since the task was very easy. Then, the criterion was sped up so participants mostly lost money and reached −6£ as the task became very hard. For the rest of the task, a linear function was applied to the RT data of the last twenty trials to set the criterion that scaled with the amount of accumulated money, if the participant was lower than 24£ in their balance, the task was easier, and if they had more it was harder and the difference in criterion was proportional to the difference in money. This procedure ensured that participants received approximately the same total of money after the experiment and experienced the different contingencies sufficiently frequently. After finishing 180 trials of the task, the participants were asked to indicate which planets predicted which outcome, by showing them the planets consecutively and asking whether it was possible (1/ “I could win money.” 2/ “I could NOT lose or win money.” 3/ “I could lose money.”).

Similar as in the learning task, participants could keep track of their progress on a bar on the right-hand side. Moreover, they could also see in real-time what their current balance was at the bottom of the screen.

### Retrieval task

Similar to the learning task, a bar on the right side tracked the progress of the task. Furthermore, a bar on the bottom side, the “Map Quality Monitor”, revealed the participant’s performance during the task, the bar was updated after each feedback slide. Participants obtained more stars by being correct for the distance 1 compared to the other distance (2, 3, 4 and plus) as they were less numerous. This feedback system used stars similar to the learning task and was used for the two first parts (0 stars – Try harder, 1 star – Well done, 2 stars – Superb, 3 stars – Amazing). The feedback appeared every 9 trials for the first part (corresponding to the pictures associations), and every 3 trials for the second part (corresponding to the planet names).

### Control tasks

The PANAS contains 10 positive (e.g.; interested) and 10 negatives adjective (e.g.; scared) describing the participant’s current mood with a scale ranging from 1= “not at all” to 5= “very much”. The SSS asks the participant to choose from 1 = “Feeling active, vital, alert, or wide awake” to 8 = “Asleep”. During the PVT, participants were facing a red counter on a black screen and were asked to press the spacebar as soon as the clock would start to count. Pressing the space would stop the counter and show to the participant their reaction time. Then, the inverse of the mean of their reaction speed (i.e.: 1/mean RT) was calculated for each phase. The word generation instructed to write as many words as possible in two minutes for either letter cues (p or m) or a category (occupations or hobbies).

### Pilot testing

Before obtaining the final paradigm described above, 3 pilot participants were run only on the learning task and another 6 pilot participants underwent the complete procedure of one experimental session. All the gamification aspects were added during the development of the paradigm in tight dialogue between the researchers and the participants to ensure high motivation and clear task instructions. The analysis strategy of the results is based mainly on the experience made during the pilot study.

An analysis of the pilot data suggested that the task could be learnt at a 70-80% accuracy level and retrieved with an overall hit rate of 60% (which was significantly higher than the overall false alarm rate). Although the learning and the retrieval accuracy were encouraging regarding feasibility of the paradigm, we decided to change the retrieval task in two distinct ways to improve on this. Initially, participants were asked if the two nodes presented were neighbours (i.e., “Was it possible to teleport between these two planets” [during learning]) and the participant could answer how sure they were from a scale from 1 to 4 (1/ “Yes, I am sure this was possible.” 2/ “Yes, but I am not sure this was possible.” 3/ “No, but I am not sure this was impossible.” 4/ “No, I am sure this was impossible.”). We changed this to the procedure asking for a distance judgement described above that allows a finer grained analysis of the graph representation. The same confidence scale (1/ “Yes, I am sure it was.” 2/ “Yes, but I am not sure it was.” 3/ “No, but I am not sure it was not.” 4/ “No, I am sure it was not.”) was initially used for the second part of retrieval that, where we showed planet name pairs and asked whether they belonged together (i.e., “Was this planet named…?”). We changed this design to the four alternatives forced choice procedure described above that reduced the amount of items to 27, as participants had complained about the length of the original task (729 items).

### Data reduction and statistical analysis

The retrieval data were analysed by calculating the hit rates (the number of times pairs of a certain distance were correctly identified,divided by all the possible pairs of this distance) for distance 1 to distance 4. Then, a repeated-measure 2 x 4 ANOVA (sum of squares type III) with the factors sleep/wake and the 4 distances was done. In addition, a paired t-test was performed between the sleep and the wake condition on the general retrieval performance calculated by taking the mean over the four distances for each participant.

Data from the learning task was reduced by calculating the mean of the accuracy for each third of the task, i.e., the mean of routes 1-27, 28-54 and 55-81 for each participant and each retention interval. Then, a repeated-measure 2 x 3 ANOVA (sum of squares type III) with the factors sleep/wake and task third was performed to assess the learning across the task. A paired t-test was completed between the sleep and the wake condition on the last third to further assure comparability. Using the method described below, the learning performance for each distance was used to create a mean for both retention intervals to get the general learning performance. Then, a paired t-test was completed between the two conditions.

To obtain the learning results for each distance, we weighed each edge of the graph to take into account all the learning data. Therefore, for distance 1, the weight for a transition depended on its position in the learning task. For example, if participants saw a transition 8 times, the weight for the first time they saw it would be 1/8, second time would be 2/8 and last time would be 8/8. Next, the weights were multiplied by the accuracy (1/0), summed together and divided by the sum of all the weights in order to give a single accuracy value between 1 and 0 for each transition. This ensured that instances closer to the end of the task were weighted more for learning accuracy than instances at the beginning. For distances 2 to 4, the graph structure was used to calculate performance between two given nodes. Specifically, the weighted accuracies of the distance 1 edges along the path between the two nodes were multiplied. Finally, the weighted performance for each distance was normalized separately by using the mean and standard deviation calculated with the data from both sleep and wake. The retention measure between learning and retrieval was calculated by deducting the normalized learning data from the normalized retrieval performance for each distance (see also Supplementary Figure 5).

Using the retention measure data, a paired t-test was performed between the sleep and the wake condition on the general performance by averaging over the distances. Additionally, a repeated-measure 2 x 4 ANOVA (sum of squares type III) with the factors sleep/wake and the 4 distances was calculated. Single paired t-tests were performed between the retention intervals for each distance. In addition, to account for the order of the distances, a regression line was created for each interval. Then, a linear model was applied to compare if there was a difference of intercept or slopes between the two regression lines.

The retention measures for each centrality (i.e. degree and closeness) were computed by averaging the retention measure data between the nodes with the centrality of interest (e.g. all nodes with a value of degree centrality of 3) and all the other nodes of the graph connected to them from distance 1 to 4 (e.g. for distance 1, it includes the neighbouring nodes of each node with a value of degree centrality of 3. Then, the nodes connected with a distance of 2, 3 and 4). Next, a repeated-measure 2 x 3 ANOVA (sum of squares type III) with the factors sleep/wake and the centrality measure was performed. In addition, paired t-tests were performed between the retention intervals for each centrality. Moreover, the method using regressions lines as described for distances above was used for both centrality measures.

The effect of reinforcement was calculated using a similar method to the centrality analysis. For each reinforced node (i.e. reward, punishment and neutral), the retention measure was calculated using the mean between the weighted edge accuracies of the reinforced node of interest (e.g. reward node) and the nodes being on the same arm of the graph. For example, if we use the node as numbered in Figure 2, the nodes we would compare to the reward node would be 6, 16, 17, 18, 25, 26 and 27 as we are excluding the central nodes (i.e. 1, 2 and 3). Then, one repeated-measure 2 x 3 ANOVA (sum of squares type III) with the factors sleep/wake and the 3 reinforced arms was performed. Finally, several paired t-tests were done between the sleep and the wake condition for each of the reinforced nodes.

The retrieval of planet-name associations was analyzed by summing the correct answers and dividing them by the number of items for each condition. A paired t-test was then performed to compare the two retention intervals.

Regarding the control tests, pen and paper data (SSS, PANAS and Word generation) were transferred into excel files and scored according to the test instructions. A single missing value in the PANAS was replaced by the mean of the items that were not missing within that scale for the participant. For the word generation task, sum scores for the letter and category cues were added together to create an overall score of retention performance for each retrieval session. The RT data from the PVT were transformed to reaction speed by dividing one by the RT for each trial. Then, the trials were averaged for each condition and participant. For all the control tasks paired t-tests were applied to compare sleep and wake.

The NSQ and SBSOD were scored according to the instructions. A unique missing value resulted in not taking the specific question into account and dividing the final score with the number of questions minus the number of questions missing. Thus, the NSQ gave a mapping tendency score from −14 to 14 and the SBSOD a score from 1 to 7 where the higher the score, the better one’s perceived sense of direction. A first pearson correlation compared the scores for both navigation scores. Then, pearson correlations were done between the NSQ scores and both intervals for learning, retrieval and retention measure data.

---

**Supplementary Figure 1.**
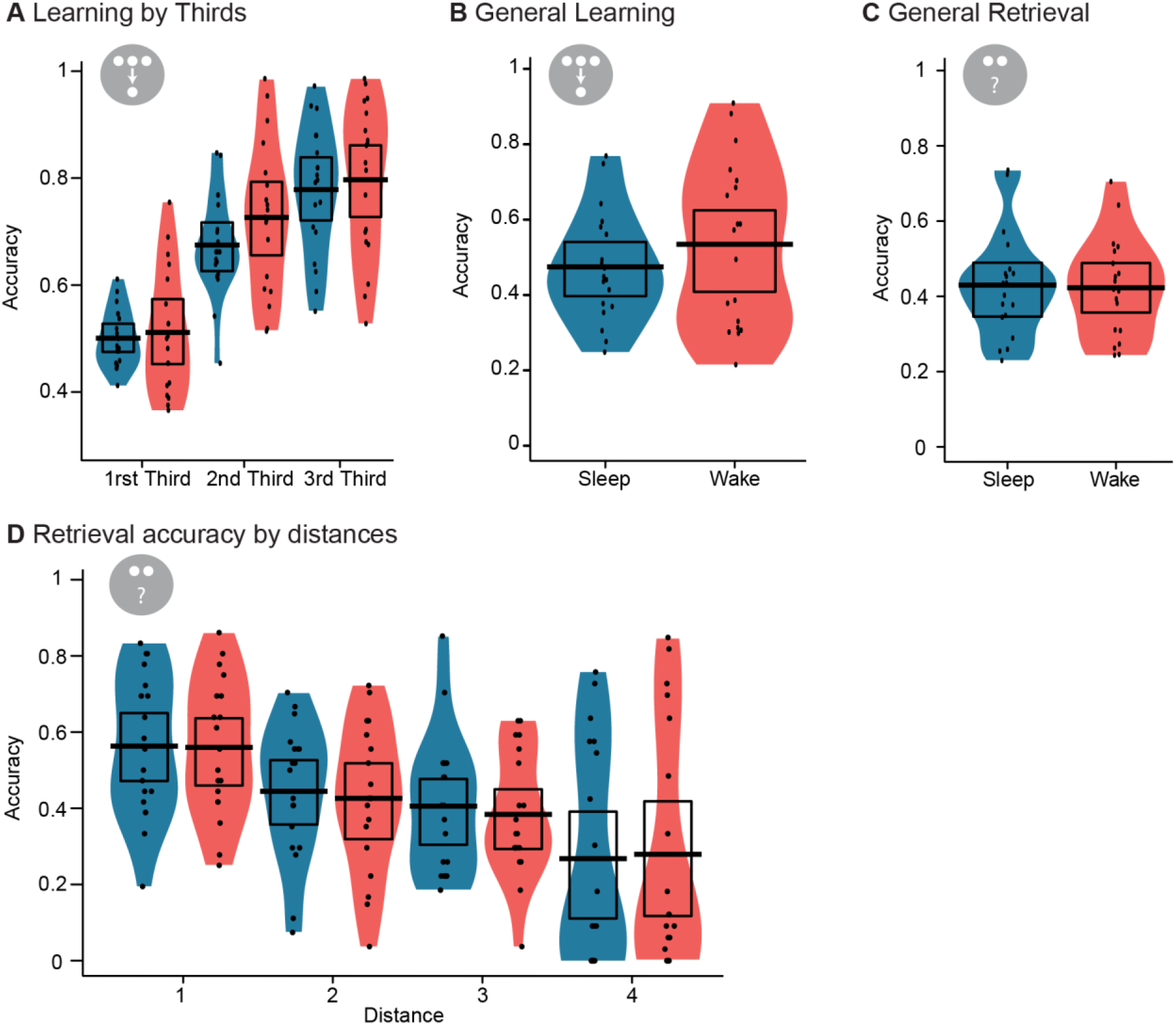
Learning and retrieval performance. A) Learning performance (proportion correct) during the learning task for the first, second and last third of the task (27 routes each) and B) for the whole task. C) Retrieval performance for the whole task. D) Retrieval performance for the different distances. The black dots, bar and rectangle represent the individual performances, the mean and the 95% of a Bayesian highest density interval, respectively. The coloured shape around shows the smoothed density.

**Supplementary Figure 2.**
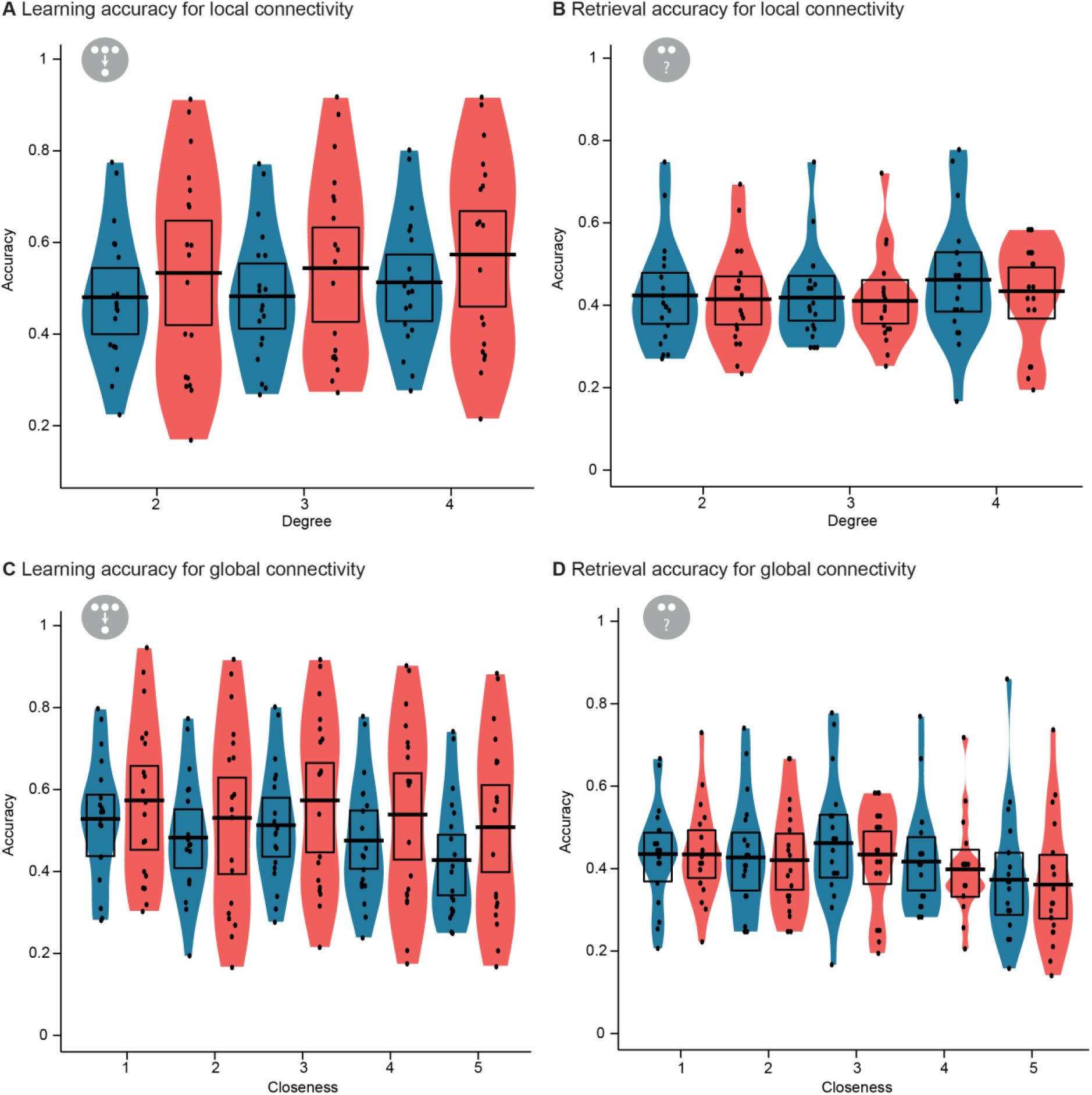
Learning and retrieval performance for centrality measures. A) Learning and B) retrieval performance for different levels of degree centrality (local connectivity). C) Learning and (d) retrieval performance for different levels of closeness centrality (global connectivity). The black dots, bar and rectangle represent the individual performances, the mean and the 95% of a Bayesian highest density interval, respectively. The coloured shape around shows the smoothed density.

**Supplementary Figure 3.**
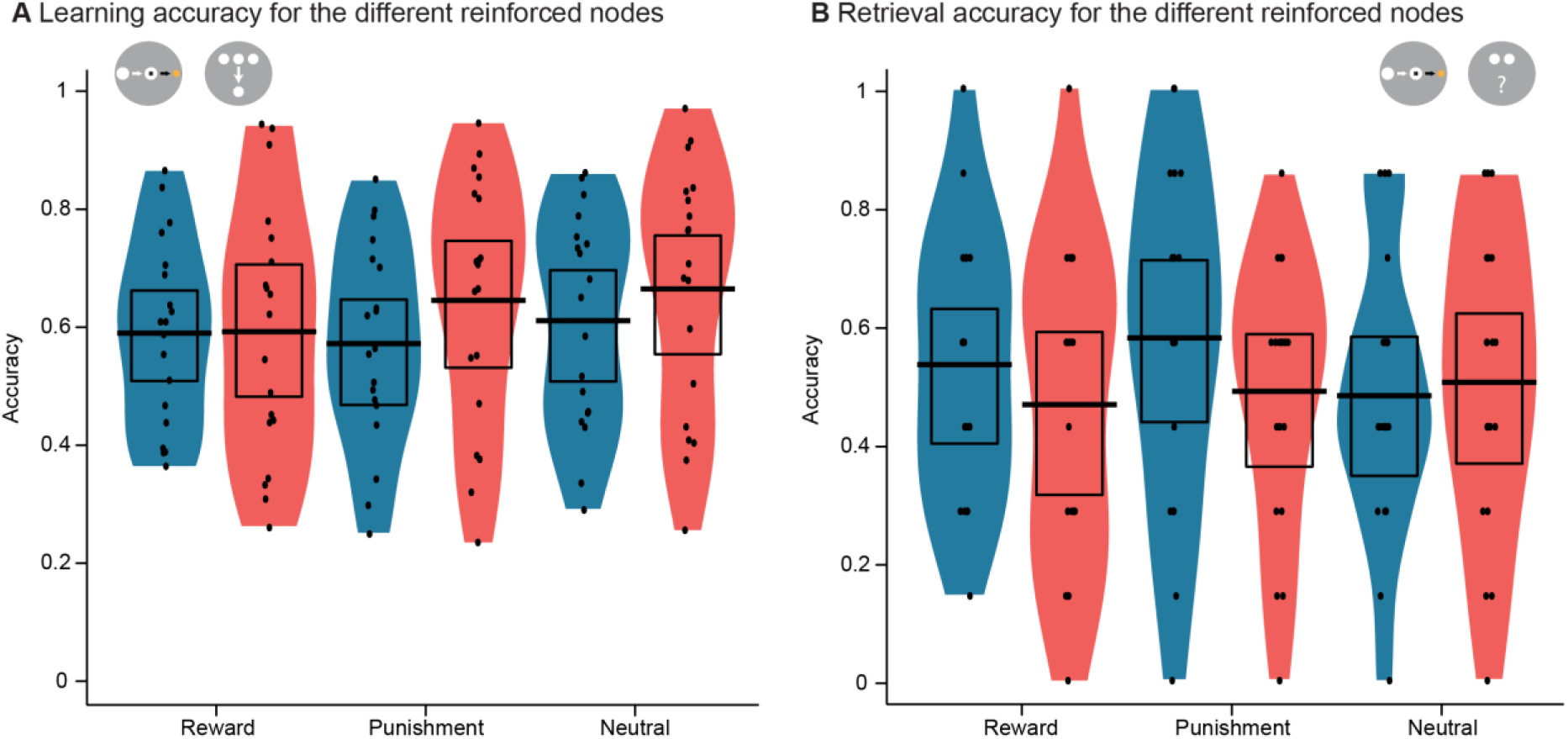
Learning and retrieval performance for the reinforced nodes. A) Learning and B) retrieval performance across the three reinforced nodes. The black dots, bar and rectangle represent the individual performances, the mean and the 95% of a Bayesian highest density interval, respectively. The coloured shape around shows the smoothed density.

**Supplementary Figure 4.**
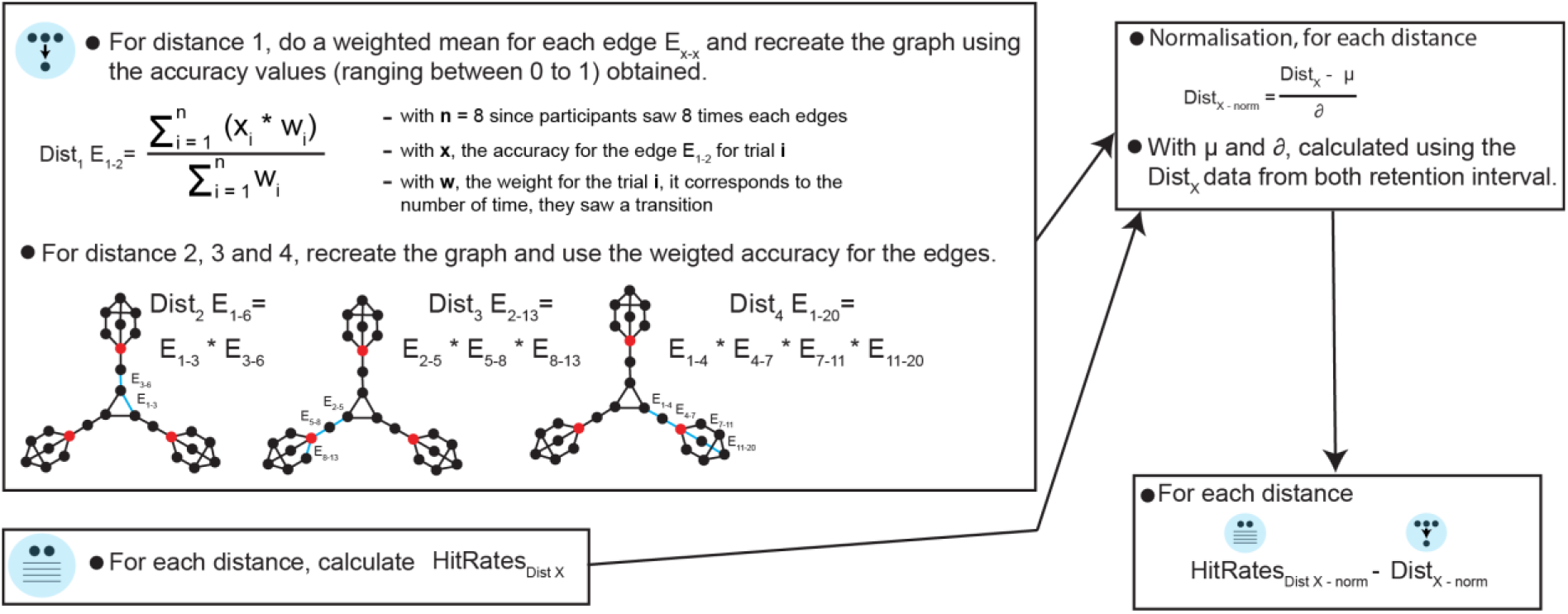
Schematic explaining the analysis to obtain the retention measure.

